# Assessment of pharmacogenomic agreement

**DOI:** 10.1101/048470

**Authors:** Zhaleh Safikhani, Nehme El-Hachem, Rene Quevedo, Petr Smirnov, Anna Goldenberg, Nicolai Juul Birkbak, Christopher E. Mason, Christos Hatzis, Leming Shi, Hugo JWL Aerts, John Quackenbush, Benjamin Haibe-Kains

## Abstract

In 2013 we published an analysis demonstrating that drug response data and gene-drug associations reported in two independent large-scale pharmacogenomic screens, Genomics of Drug Sensitivity in Cancer^1^ (GDSC) and Cancer Cell Line Encyclopedia^2^ (CCLE), were inconsistent^3^. The GDSC and CCLE investigators recently reported that their respective studies exhibit reasonable agreement and yield similar molecular predictors of drug response^4^, seemingly contradicting our previous findings^3^. Reanalyzing the authors’ published methods and results, we found that their analysis failed to account for variability in the genomic data and more importantly compared different drug sensitivity measures from each study, which substantially deviate from our more stringent consistency assessment. Our comparison of the most updated genomic and pharmacological data from the GDSC and CCLE confirms our published findings that the measures of drug response reported by these two groups are not consistent^5^. We believe that a principled approach to assess the reproducibility of drug sensitivity predictors is necessary before envisioning their translation into clinical settings.

## Introduction

Pharmacogenomic studies correlate genomic profiles and sensitivity to drug exposure in a collection of samples to identify molecular predictors of drug response. The success of validation of such predictors depends on the level of noise both in the pharmacological and genomic data. The groundbreaking release of the Genomics of Drug Sensitivity in Cancer^1^ (GDSC) and Cancer Cell Line Encyclopedia^2^ (CCLE) datasets enables the assessment of pharmacogenomic data consistency, a necessary requirement for developing robust drug sensitivity predictors. Below we briefly describe the fundamental analytical differences between our initial comparative study^3^ and the recent assessment of pharmacogenomic agreement published by the GDSC and CCLE investigators^4^.

### Comparison of drug sensitivity predictors

Given the complexity and high dimensionality of pharmacogenomic data, the development of drug sensitivity predictors is prone to overfitting and requires careful validation. In this context, one would expect the most significant predictors derived in GDSC to accurately predict drug response in CCLE and *vice versa*. This will be the case if both studies independently produce consistent measures of both genomic profiles and drug response for each cell line. In our comparative study^3^, we made direct comparison of the same measurements generated independently in both studies by taking into account the noise in both the genomic and pharmacological data (Figure 1a). By investigating the authors’ code and methods, we identified key shortcomings in their analysis protocol, which have contributed to the authors’ assertion of consistency between drug sensitivity predictors derived from GDSC and CCLE.

For their ANOVA analyses, the authors used drug activity area (1-AUC) values independently generated in GDSC and CCLE, but used the same GDSC mutation data across the two different datasets (Figure 1b; see Methods). By using the same mutation calls for both GDSC and CCLE, the authors have disregarded the noise in the molecular profiles, while creating an information leak between the two studies. For their ElasticNet analysis, the authors followed a similar design by reusing the CCLE genomic data across the two datasets, but comparing different drug sensitivity measures that are IC_50_ in GDSC vs. AUC in CCLE (Figure 1c; see Methods).

**Figure 1:**
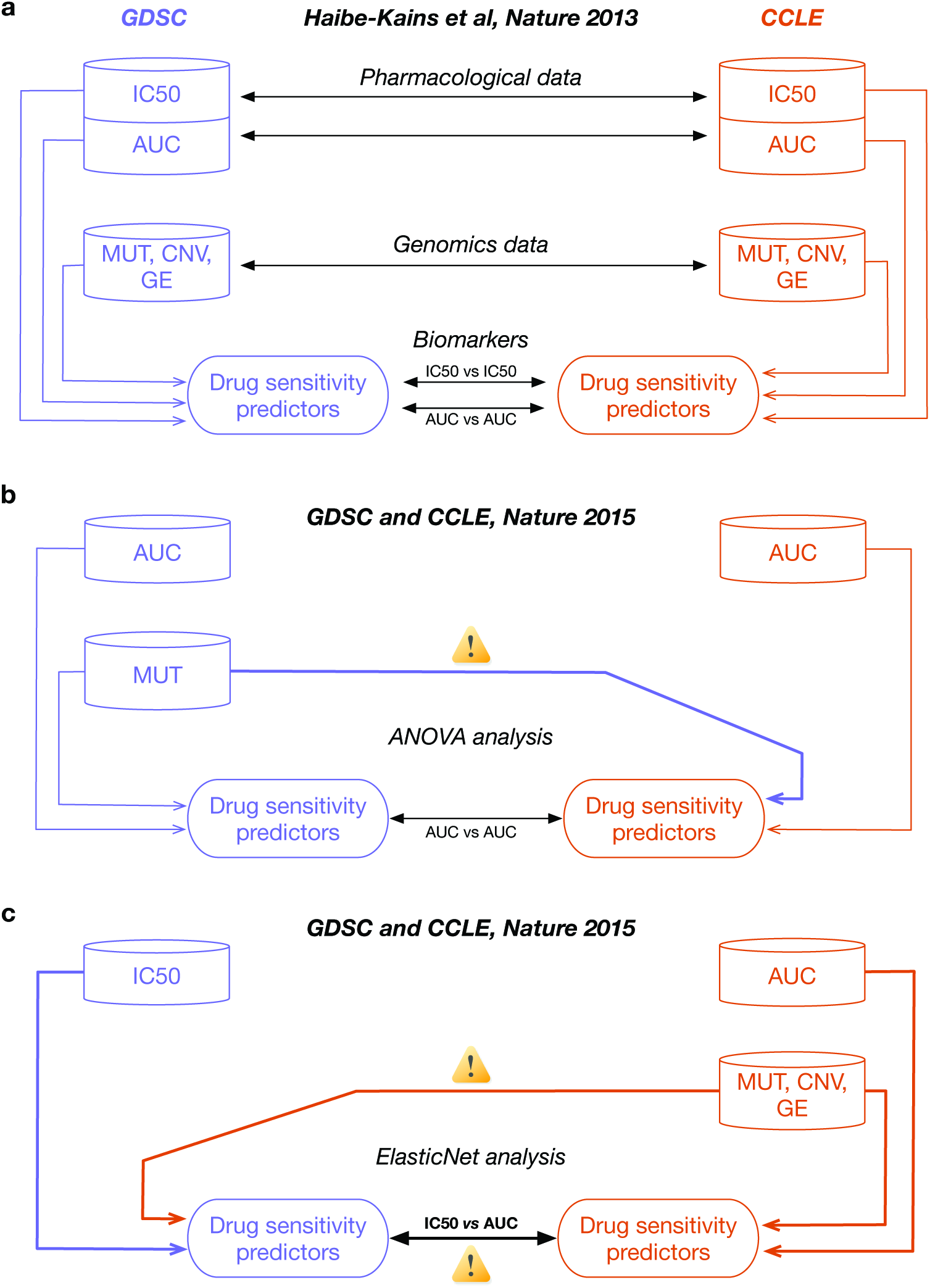
Analysis designs used to compare pharmacogenomic studies. (a) Analysis design used in our comparative study (Haibe-kains et al, Nature 2013) where each data generated by GDSC and CCLE are independently compared to avoid information leak and biased assessment of consistency. (b) Analysis design used by the GDSC and CCLE investigators for their ANOVA analysis where the mutation data generated with GDSC were duplicated for use in the CCLE study. (c) Analysis design for the ElasticNet analysis where the molecular profiles from CCLE where duplicated in the GDSC study and the GDSC IC_50_ were compared to CCLE AUC data. Differences between our analysis design and those used by the GDSC and CCLE investigators are indicated by yellow signs with exclamation mark symbol.

We are puzzled by the seemingly arbitrary choices of analytical design made by the authors, which raises the question as to whether the use of different genomic data and drug sensitivity measures would yield the same level of agreement. Moreover, by ignoring the (inevitable) noise and biological variation in the genomic data, the authors’ analyses is likely to yield over-optimistic estimates of data consistency, as opposed to our more stringent analysis design^3^.

### What constitutes agreement?

In examining correlation, there is no universally accepted standard for what constitutes agreement. However, the FDA/MAQC consortium guidelines define good correlation for inter-laboratory reproducibility^6–9^ to be ≥0.8. The authors of the present study used two measures of correlation, Pearson correlation (ρ) and Cohen’s kappa (ϰ) coefficients, but never clearly defined *a priori* thresholds for consistency, instead referring to ρ>0.5 as “reasonable consistency” in their discussion. Of the 15 drugs that were compared, their analysis found only two (13%) with ρ>0.6 for AUC and three (20%) above that threshold for IC_50_. This raises the question whether ρ~0.5-0.6 for one third of the compared drugs should be considered as “good agreement.” If one applies the FDA/MAQC criterion, only one drug (nilotinib) passes the threshold for consistency.

Similarly, the authors referred to the results of their new Waterfall analysis as reflective of “high consistency,” even though only 40% of drugs had a ϰ≥0.4, with five drugs yielding moderate agreement and only one drug (lapatinib) yielding substantial agreement according to the accepted standards^10^. Based on these results, the authors concluded that 67% of the evaluable compounds showed reasonable pharmacological agreement, which is misleading as only 8/15 (53%) and 6/15 (40%) drugs yielded ρ>0.5 for IC_50_ and AUC, respectively. Taking the union of consistency tests is bad practice; adding more sensitivity measures (even at random) would ultimately bring the union to 100% without providing objective evidence of actual data agreement.

### Sources of inconsistency in pharmacological data

The authors acknowledged that the consistency of pharmacological data is not perfect due to the methodological differences between protocols used by CCLE and GDSC, further stating that standardization will certainly improve correlation metrics. To test this important assertion, the authors could have analyzed the replicated experiments performed by the GDSC using identical protocols to screen camptothecin and AZD6482 against the same panel of cell lines at the Wellcome Trust Sanger Institute and the Massachusetts General Hospital.

Our re-analyses^3,5^ of drug sensitivity data from these drugs found a correlation between GDSC sites on par with the correlations observed between GDSC and CCLE (ρ=0.57 and 0.39 for camptothecin and AZD6482, respectively; Extended Data Figure 1a,b). These results suggest that intrinsic technical and biological noise of pharmacological assays is likely to play a major role in the lack of reproducibility observed in high-throughput pharmacogenomic studies, which cannot be attributed solely to the use of different experimental protocols

## Conclusions

We agree with the authors that their and our observations “[…] *raise important questions for the field about how best to perform comparisons of large-scale data sets*, *evaluate the robustness of such studies*, *and interpret their analytical outputs*.” We believe that a principled approach using objective measures of consistency and an appropriate analysis strategy for assessing the independent datasets is essential. An investigation of both the methods described in the manuscript and the software code used by the authors to perform their analysis^4^ identified fundamental differences in analysis design compared to our previous published study^3^. By taking into account variations in both the pharmacological and genomic data, our assessment of pharmacogenomic agreement is more stringent and closer to the translation of drug sensitivity predictors in preclinical and clinical settings, where zeronoise genomic information cannot be expected.

Our stringent reanalysis of the most updated data from the GDSC and CCLE confirms our 2013 finding that the measures of drug response reported by these two groups are not consistent and have not improved substantially as the groups have continued generating data since 2012^5^. While the authors make arguments suggesting consistency, it is difficult to imagine using these post hoc methods to drive discovery or precision medicine applications.

The observed inconsistency between early microarray gene expression studies served as a rallying cry for the field, leading to an improvement and standardization of experimental and analytical protocols, resulting in the agreement we see between studies published today. We are looking forward to the establishment of new standards for largescale pharmacogenomic studies to realize the full potential of these valuable data for precision medicine.

## Methods

### The authors’ software source code

As the authors’ source code, we refer to the ‘CCLE.GDSC.compare’ (version 1.0.4 from December 18, 2015) and DRANOVA (version 1.0 from October 21, 2014) R packages available from http://www.broadinstitute.org/ccle/Rpackage/.

### ANOVA analysis

In the authors’ ANOVA analyses, identical mutation data were used for both GDSC and CCLE studies as can be seen in the authors’ analysis code in lines 20, 25-35 of CCLE.GDSC.compare::plotFig2A_biomarkers.R.

### ElasticNet (EN) analysis

In their EN analyses, the authors compared different drug sensitivity measures, using IC_50_ in GDSC and AUC in CCLE, as described in the Supplementary Data 5 and stated in the Methods section of their published study:

> *“Since the IC50 is not reported in CCLE when it exceeds the tested range of 8 μM, we used the activity area for the regression as in the original CCLE publication. We also used the values considered to be the best in the original GDSC study: the interpolated log(IC50) values.”*

This was confirmed by looking at the authors’ analysis code, lines 83 and 102 of CCLE.GDSC.compare::ENcode/prepData.R. Moreover, identical genomic data were used for both GDSC and CCLE studies, as described the Methods section of the published study:

> *“In order to compare features between the two studies, we used the same genomic data set (CCLE).”*

This was confirmed by looking at the authors’ analysis code, lines 17, 38, 51, and 70 of CCLE.GDSC.compare::ENcode/genomic.data.R, and lines 10-11 of CCLE.GDSC.compare::plotFigS6_ENFeatureVsExpected.R.

### Statistical analysis

All analyses were performed using the most updated version of the GDSC and CCLE pharmacogenomic data based on our *PharmacoGx* package^11^ (version 1.1.4).

## Supplementary Files

*Supplementary File 1*. Supplementary Information, including additional comments and supplemental methods.

*Supplementary File 2*. Documented R code used to generate all the analysis results and figures.

**Extended Data Figure 1:**
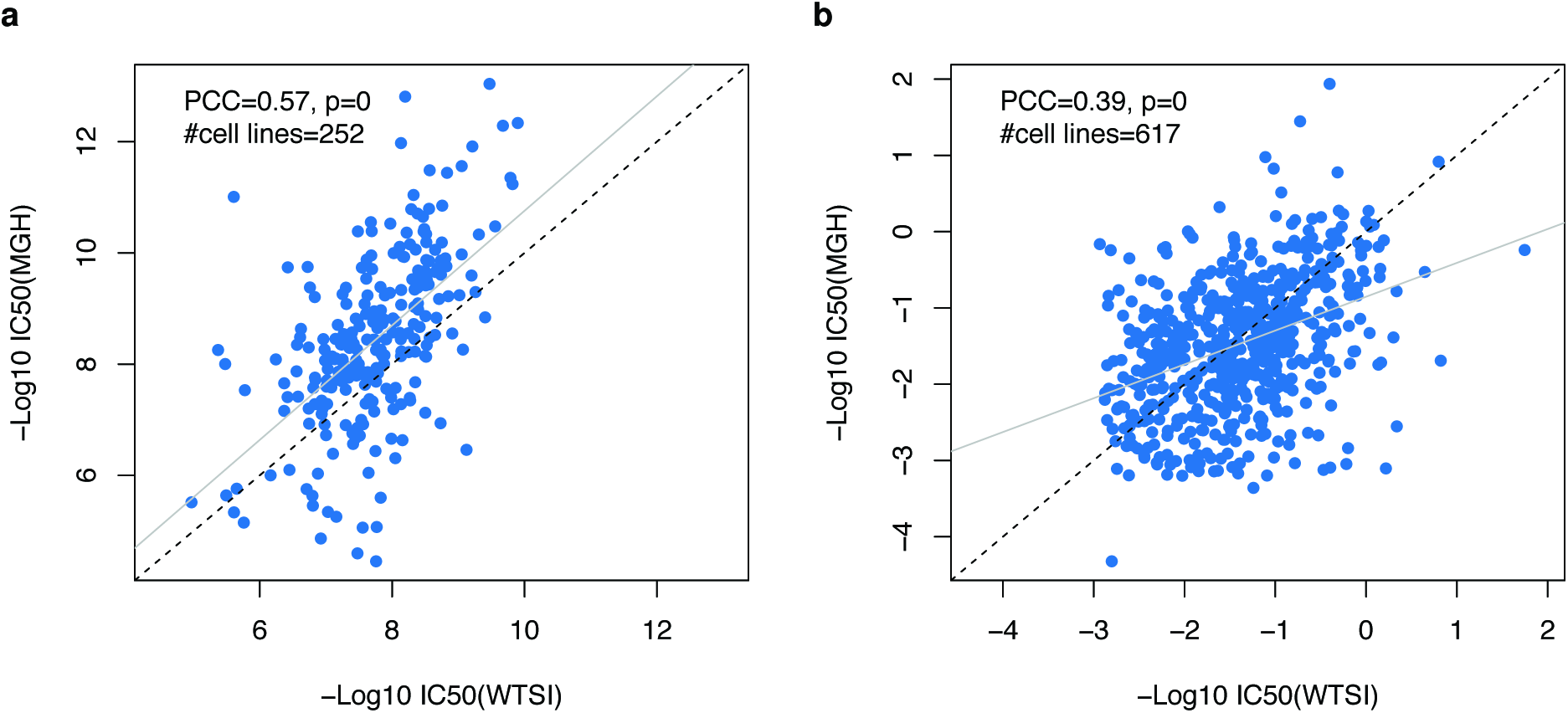
Consistency of sensitivity profiles between replicated experiments across GDSC sites: (a) Camptothecin and (b) AZD6482. PCC: Pearson correlation coefficient.

**Extended Data Figure 2:**
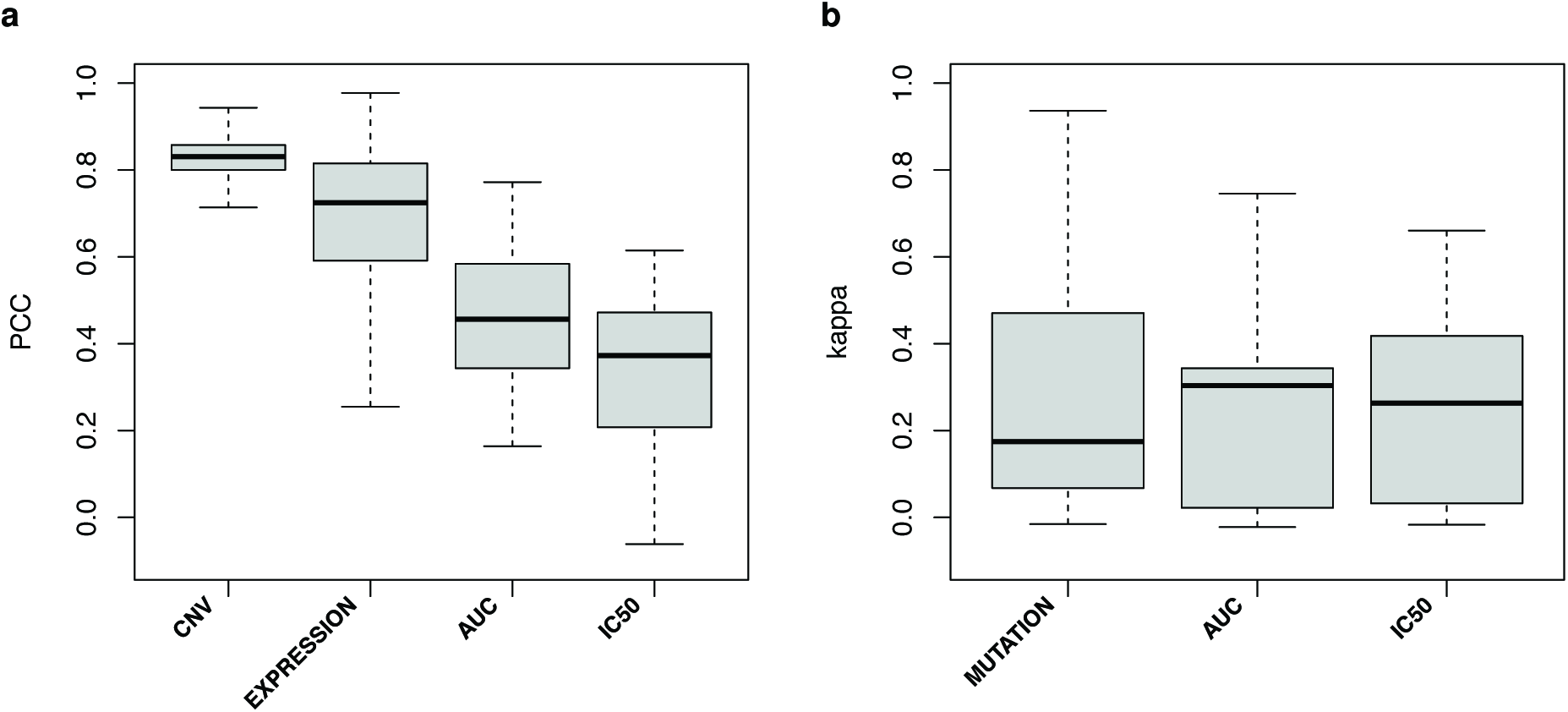
Consistency of molecular profiles between GDSC and CCLE: (a) Continuous values for gene copy number ratio (CNV), gene expression (EXPRESSION), AUC and IC_50_ and (b) for binary values for presence/absence of mutations (MUTATION) and insensitive/sensitive calls based on AUC and IC_50_ values. PCC: Pearson correlation coefficient; Kappa: Cohen’s Kappa coefficient.

## SUPPLEMENTARY INFORMATION

**Assessment of pharmacogenomic agreement**

### Additional comments

#### Which pharmacological drug response data should one use?

The first GDSC and CCLE studies were published in 2012 and the investigators of both studies have continued to generate data and to release them publicly. One would imagine that any comparative study would use the most current versions of the data. However, the authors of the reanalysis used an old release of the GDSC (July 2012) and CCLE (February 2012) pharmacological data, resulting in the use of outdated IC50 values, as well as missing approximately 400 new drug sensitivity measurements for the 15 drugs screened both in GDSC and CCLE. Assessing data that are three years old and which have been replaced by the very same authors with more recent data seems to be a substantial missed opportunity. It raises the question as to whether the current data would be considered to be in agreement and which data should be used for further analysis.

#### Consistency of genomic data

In their comparative study, the authors did not assess the consistency of genomic data between GDSC and CCLE. Our recent reanalysis [3] reaffirmed that consistency of gene copy number and expression data was significantly higher than for drug sensitivity data (one-sided Wilcoxon rank sum test p-value=3x10-5; Extended Data Figure 2), while mutation data exhibited poor consistency as reported previously [2]. The very high consistency of copy number data is quite remarkable and could be partly attributed to the fact that CCLE investigators used their SNP array data to compare cell line fingerprints with those of the GDSC project prior to publication and removed the discordant cases from their dataset [1].

#### The Waterfall approach

In the Methods, the authors use all cell lines to optimally identify the inflexion point in the response distribution curves. The authors stated that "This is a major difference to the Haibe-Kains et al. analysis, as that analysis only considered the cell-lines in common between the studies when generating response distribution curves." This is not correct. As can be seen in our publicly available R code, we performed the sensitivity calling (using the Waterfall approach as published in [1]) before restricting our analysis to the common cell lines, for the obvious reasons that the authors mentioned in their manuscript. See lines 308 and 424 in https://github.com/bhklab/cdrug/blob/master/CDRUG_format.R.

### Supplemental Methods

#### Pharmacogenomic data

As evidenced in the authorsâĂŹ code (lines 20 and 29 of CCLE.GDSC.compare::PreprocessData.R), they used GDSC and CCLE pharmacological data released on July 2012 and February 2012, respectively. However the GDSC released updated sets of pharmacological data (release 5) on June 2014, gene expression arrays (E-MTAB-3610) and SNP arrays (EGAD00001001039) on July 2015. CCLE released updated pharmacological data on February 2015, the mutation and SNP array on October 2012, and the gene expression data, on March 2013. These updates substantially increased the overlap in genomic features between the two studies, thus providing new opportunities to investigate the consistency between GDSC and CCLE10.

#### Research reproducibility

All analyses were performed using the most updated version of the GDSC and CCLE pharmacogenomic data based on our PharmacoGx package11 (version 1.1.3). PharmacoGx provides intuitive function to download, intersect and compare large pharmacogenomics datasets. The PharmacoSet for the GDSC and CCLE datasets are available from pmgenomics.ca/bhklab/sites/default/files/downloads/ using the downloadPSet() function. The code and the data used to generate all the results and figures is available as Supplementary File 2.

#### Acronyms

ABC: Area between the curves
AE: ArrayExpress by the European Bioinformatics Institute
AUC: Area upper the dose response curve
CCLE: The Cancer Cell Line Encyclopedia initiated by the Broad Institute of MIT and Harvard
EN: ElasticNet analysis
GDSC: The Cancer Genome Project initiated by the Wellcome Trust Sanger Institute
IC_50_: Concentration at which the drug inhibited 50% of the maximum cellular growth
KAPPA: Cohen’s *k* coefficient of agreement
PCC: Pearson product-moment correlation coefficient
SNP: Single nucleotide polymorphism

#### Session Information

- R version 3.2.3 (2015-12-10), `x86_64-apple-darwin13.4.0`
- Locale: `en_CA.UTF-8/en_CA.UTF-8/en_CA.UTF-8/C/en_CA.UTF-8/en_CA.UTF-8`
- Base packages: base, datasets, graphics, grDevices, methods, stats, utils
- Other packages: PharmacoGx 1.1.4
- Loaded via a namespace (and not attached): Biobase 2.28.0, BiocGenerics 0.14.0, bitops 1.0-6, caTools 1.17.1, cluster 2.0.3, digest 0.6.8, downloader 0.4, gdata 2.17.0, gplots 2.17.0, gtools 3.5.0, igraph 1.0.1, KernSmooth 2.23-15, limma 3.24.15, magicaxis 1.9.4, magrittr 1.5, marray 1.46.0, MASS 7.3-45, parallel 3.2.3, piano 1.8.2, plotrix 3.6-1, RColorBrewer 1.1-2, relations 0.6-6, sets 1.0-16, slam 0.1-32, sm 2.2-5.4

